# Identification of cell types, states and programs by learning gene set representations

**DOI:** 10.1101/2023.09.08.556842

**Authors:** Soroor Hediyeh-zadeh, Holly J. Whitfield, Malvika Kharbanda, Fabiola Curion, Dharmesh D. Bhuva, Fabian J. Theis, Melissa J. Davis

**Affiliations:** Bioinformatics Division, The Walter and Eliza Hall Institute of Medical Research, Parkville, VIC 3052, Australia; Department of Medical Biology, The Faculty of Medicine, Dentistry and Health Science, The University of Melbourne, Carlton, VIC 3010, Australia; Institute of Computational Biology, Department of Computational Health, Helmholtz Munich, Munich, Germany; Department of Mathematics, School of Computation, Information and Technology, Technical University of Munich, Garching, Germany; TUM School of Life Sciences Weihenstephan, Technical University of Munich, Munich, Germany; Munich Center for Machine Learning, Technical University of Munich, Garching, Germany; Department of Clinical Pathology, Faculty of Medicine, Dentistry and Health Science, The University of Melbourne, Carlton, VIC 3000, Australia; South Australian Immunogenomics Cancer Institute, Faculty of Health and Medical Sciences, The University of Adelaide, Adelaide, Australia; The University of Queensland Diamantina Institute, The University of Queensland, Brisbane, QLD 4102 Australia

## Abstract

As single cell molecular data expand, there is an increasing need for algorithms that efficiently query and prioritize gene programs, cell types and states in single-cell sequencing data, particularly in cell atlases. Here we present scDECAF, a statistical learning algorithm to identify cell types, states and programs in single-cell gene expression data using vector representation of gene sets, which improves biological interpretation by selecting a subset of most biologically relevant programs. We applied scDECAF to scRNAseq data from PBMC, Lung, Pancreas, Brain and slide-tags snRNA of human prefrontal cortex for automatic cell type annotation. We demonstrate that scDECAF can recover perturbed gene programs in Lupus PBMC cells stimulated with IFNbeta and TGFBeta-induced cells undergoing epithelial-to-mesenchymal transition. scDECAF delineates patient-specific heterogeneity in cellular programs in Ovarian Cancer data. Using a healthy PBMC reference, we apply scDECAF to a mapped query PBMC COVID-19 case-control dataset and identify multicellular programs associated with severe COVID-19. scDECAF can improve biological interpretation and complement reference mapping analysis, and provides a method for gene set and pathway analysis in single cell gene expression data.

## Introduction

The maturation of single-cell sequencing technologies has increased the capacity to generate vast amounts of single-cell data, leading to the emergence of cell atlases such as the Human Cell Atlas^1^. These atlases have opened the way to understanding cell types, disease-associated genes, and programs in which they act. The identification of cell types, states and gene programs allows researchers to decipher the mechanisms of disease, empowering the development of new drugs and therapies. Cell types and states can be identified using various computational methods^2,3,4,5^ that transfer annotations from single-cell references^6,7,8^, by means of tissue-specific cell type markers^9,10^ or classifiers^3,4^. For example, Seurat V4^6^ and SingleR^7^ leverage existing reference scRNAseq datasets for cell type annotation, whereas CellAssign^9^ and Garnett^10^ can deliver the same task with cell type markers.

Gene programs can capture coordinated differences between cell populations and represent subtle, continuous cell phenotypes that are not captured by cell type annotations. Cell-intrinsic gene programs can be identified by scoring single-cell data with gene sets that represent molecular pathways or transcriptional signatures. Scoring methods apply gene signature knowledge-bases using rank-based or factorization-based approaches^11,12,13^ to obtain per-cell scores that facilitate the interpretation of cellular phenotypes across a dataset. For example, Spectra^13^ is a Bayesian approach that can discern cell type-specific phenotypes from global programs while refining the definition of a gene set depending on the context. Alternatively, to obtain gene programs that span across cell types, recent methods like DIALOGUE^14^, aim to uncover multicellular processes in single-cell RNAseq data.

Deep learning-based solutions based on Variational Autoencoders, including VEGA^15^ and expiMap^16^, learn a disentangled lower-dimensional representation of gene expression in cells, whereby each dimension of the latent space corresponds to an input gene program. While both VEGA and expiMap are scalable approaches to quantify known gene programs in datasets with millions of cells, expiMap additionally allows for selection of a subset of gene programs that are more relevant for reconstruction of gene expression. These methods identify cell type-specific enrichment of gene programs in diseases or perturbations, however, since Variational Autoencoders are inherently data denoising and compression techniques, they are susceptible to the strength of the biological signal or effect size of the perturbation in the data and may not perform optimally in the absence of sufficiently large biological effects. Furthermore, interpretable deep learning approaches can not be applied to integrated cell atlases such as the adult lung cell atlas^17^ that contains 2.2 million cells from across multiple studies. In order to identify disease- or perturbation-specific gene programs in integrated atlases, the interpretable deep learning approaches have to be used at the train time for data integration.

In this work, we present a statistical learning approach that learns vector representations of gene sets to identify cell types and gene expression programs in single-cell gene expression data. Similar to interpretable deep learning models, our approach can learn disentangled representations of cellular gene expression profiles and select the most relevant subset of gene programs among a collection of gene sets. Unlike these approaches, our method can be applied downstream of data integration workflows to identification of cell types, states and programs in existing atlases. Our method is capable of a sparse selection of gene sets that prioritizes pathways and signatures to enhance biological interpretation of data, which is not offered by existing statistical learning approaches.

## Results

### Cell to gene-set assignment by similarity of vector representations in the vector space recovers gene programs

To interrogate similarities between the expression profiles of individual cells in a dataset and patterns encoded by a gene set, we developed scDECAF (**S**ingle-**c**ell **d**is**e**ntanglement by **ca**nonical **f**actors), which enables reference-free automated annotation of cells with either discrete labels, such as cell types and states, or continuous phenotype scores for gene expression programs (**Figure 1A**). Our method uses canonical correlation analysis (CCA)^18–19^ to construct a shared lower-dimensional space between binarised gene lists and unlabelled single-cell gene expression profiles. This lower-dimensional space provides vector representations of gene sets and gene expression profiles while simultaneously maximizing the correlation between the two. The association between individual cells and phenotype is determined based on the similarity of their representations in CCA space. Thus, scDECAF can leverage existing large knowledge bases in the interpretation of heterogeneous single cell datasets for either cell type annotation or gene signature scoring. To improve interpretability, scDECAF offers a “discovery mode” - a sparsity inducing procedure that supports the identification of relevant gene sets prior to learning the shared lower-dimensional space. This additional filtering step selects for gene sets that are most relevant for reconstruction of a lower dimensional embedding of cells, using a multi-response penalized linear regression model^20^ (See **Materials and Methods**). This is a principled, data-driven alternative to manual gene set selection, which can introduce unwanted bias.

**Figure 1:**
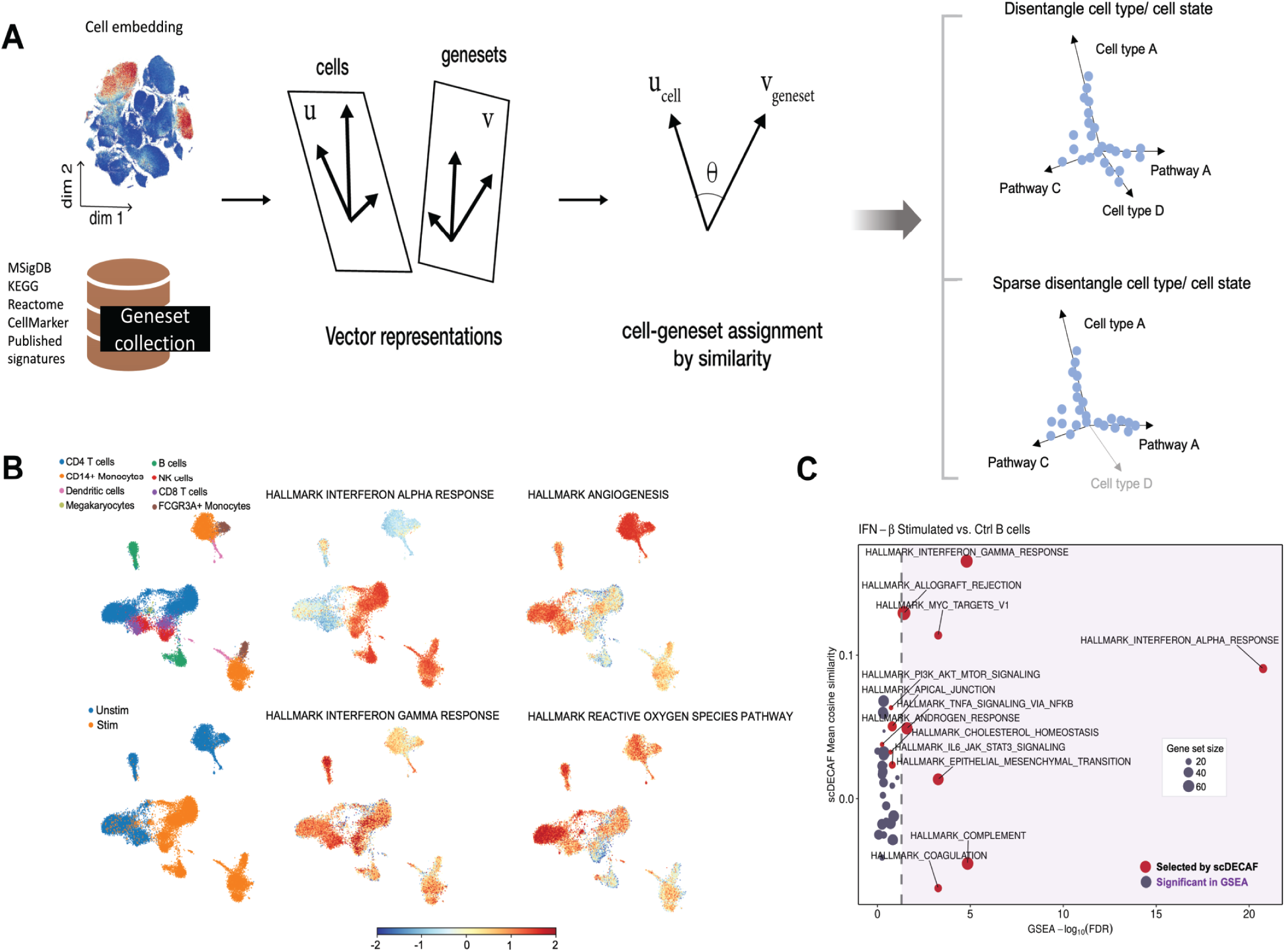
scDECAF quantifies cell states and gene programs by learning vector representations of cells and gene sets. **A.** scDECAF uses Canonical Correlation Analysis (CCA) to learn a lower-dimensional space shared between single cell gene expression profiles and gene sets. In this space, cells and gene sets are represented as vectors and there is an optimal correlation between gene expression profiles and patterns specified by the gene sets. The proximity between the vector representation of the cells and gene sets is used for cell type annotation, or to score gene programs and transcriptional signatures. Additionally, the learned space can be sparsified, which allows the selection of most biologically relevant gene set terms. **B.** The activity score of most biologically relevant Hallmark gene sets selected and quantified by scDECAF in a IFN-*β* stimulated lupus PBMC dataset. **C.** Comparison of GSEA and scDECAF in identifying Hallmark gene sets enriched in IFN-*β* stimulated B cells compared to unstimulated B cells.

We applied scDECAF to PBMC cells from individuals with Systemic Lupus Erythematosus (SLE) perturbed by IFNβ stimulation^21^ and Hallmark gene sets^22^. scDECAF correctly identified IFNα response, which is known to be induced by IFNβ^21^, in stimulated cells (**Figure 1B**). scDECAF additionally identified IFNγ response in cells of lymphoid lineage in both control and interferon stimulated cells, consistent with established role of IFNγ signaling in SLE^23^ and its specificity to the lymphoid lineage^24^. Reduced angiogenesis^25^ and oxygen reactive species pathway activity^26^ in interferon-stimulated were also identified by scDECAF, and are likewise supported by literature^25,26^ (**Figure 1B**). When compared to standard GSEA (**Figure 1C**), many Hallmark gene sets identified through the sparse gene program selection procedure of scDECAF were also statistically enriched in interferon-stimulated B cells. However, scDECAF additionally identified a number of gene programs that were not detected by standard GSEA, namely the PI3k-AKT-mTOR signaling program, which is known to be elevated in human SLE B cells^27^ . Overall these results suggest that scDECAF can recover well-studied gene programs and identify new programs not detected by conventional GSEA analysis.

### Cell to marker gene-set assignment by vector representation for cell type annotation

To benchmark scDECAF for cell type annotation, we systematically compared scDECAF to a broad range of cell type annotation tools, including marker list-based methods and those that require reference single cell gene expression datasets. We assessed the performance using published annotations as ground truth. To run scDECAF, we obtained the tissue-specific cell type markers from PanglaoDB^28^ or CellMarker^29^, which are single cell-specific cell marker knowledge bases.

We first compared scDECAF to four reference-based methods (SingleR^7^, Scmap^8^, Garnett^10^ and Seurat v4^6^) to annotate sorted PBMC cells from healthy donors^30^ (**Figure 2A, C**) and fine-grade, postnatal mouse cells from dentate gyrus^31^ . We used independent data for query and reference datasets (See **Materials and Methods**). To ensure comparable ground truths between methods that require an annotated reference scRNAseq dataset (SingleR, Scmap, Seurat v4, Garnett) and those that use input gene lists (Garnett, scDECAF), we harmonized cell type labels between the different resources by manual annotation (**Supplementary Table 3 and 6**). All methods were run on default settings and compared using F1 score (**Figure 2**).

**Figure 2:**
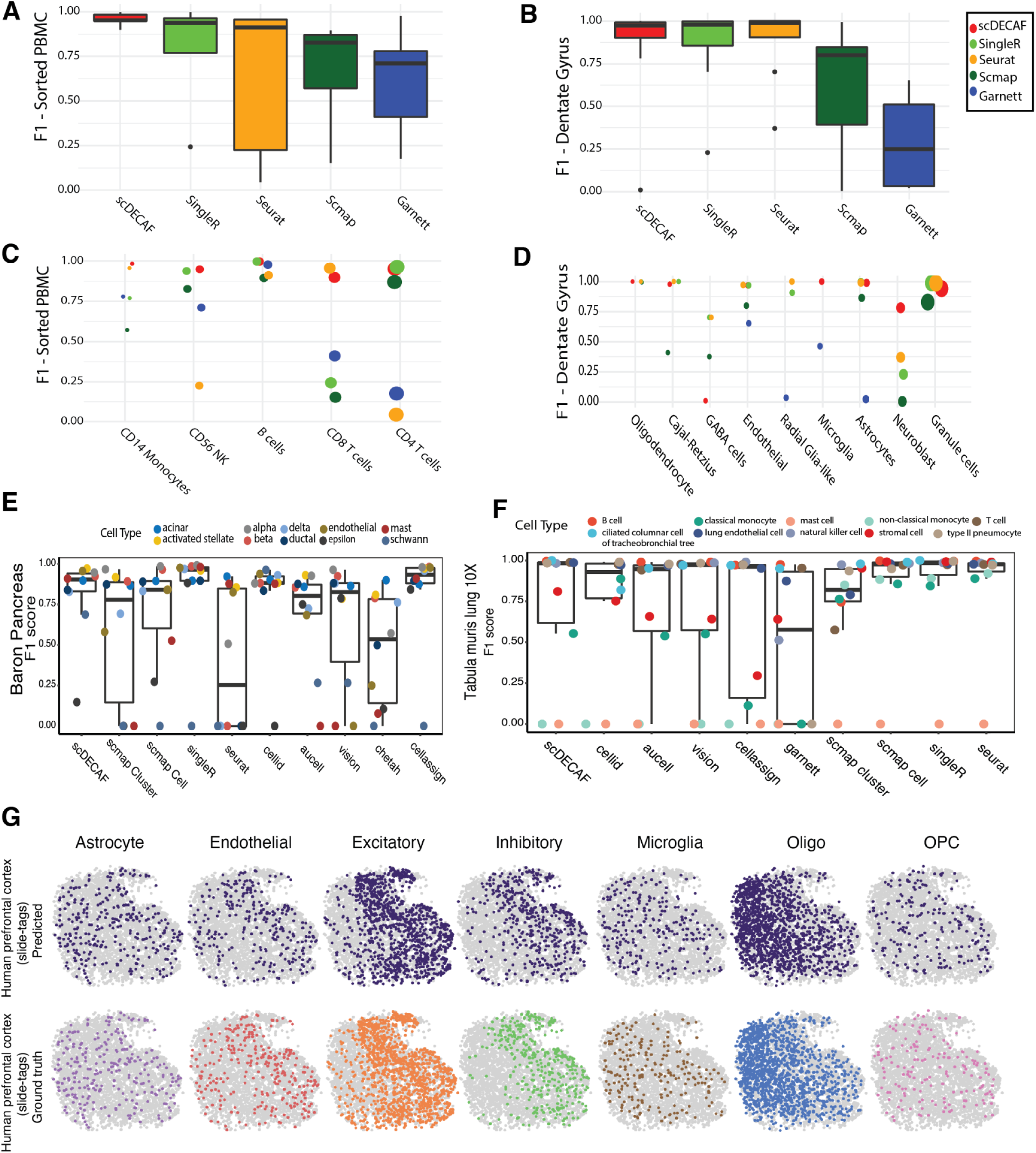
The performance of scDECAF relative to cell type annotation and gene set scoring methods. Comparison of cell type annotation accuracy in a diverse range of datasets. Per cell type F1-score is compared between methods in PBMC **(A,C)**, Dentate Gyrus **(B,D)**, Pancreas **(E)** and Lung **(F)** datasets. scDECAF cell type predictions (top row) versus marker-based annotation of cell types (bottom row) in a spatial snRNA of human prefrontal cortex measured by slide-tags **(G)**. The size of the points in **C** and **D** represent the population size for the cell type. Higher F1-score indicates better performance.

We observed a higher median F1 score across the range of cell types for scDECAF annotations relative to reference-based methods in the PBMC dataset (**Figure 2A**). The subtle differences between some lymphocyte populations, such as CD8 and CD4 T cells, can be difficult to discern and may explain the varied performance on the PBMC dataset across methods. While scDECAF outperformed scmap and Garnett in the Dentate Gyrus dataset (**Figure 2B**), the median F1 score for scDECAF was comparable to singleR and Seurat v4 owing to the failed annotation of GABA cells. Taking a closer look at the F1 score per cell type, scDECAF maintained a high accuracy across all cell types in the sorted PBMC dataset (**Figure 2C**), and across all but GABA cells in the Dentate Gyrus dataset (**Figure 2D**). We postulated that this observation could be explained by either the poor quality of marker list used to annotate this population of cells (i.e. marker list does not contain sufficiently relevant gene markers to characterize GABA cells in postnatal neurogenesis), or a high degree of similarity between marker lists for cell types present in the dataset that are different, but have very similar transcriptional profiles (i.e. marker genes for the cell types are highly overlapping, and genes are similarly expressed in two or more cell types). We, therefore, extended the evaluations to datasets with more diverse cell states.

To further investigate the robustness of our method in annotating rare cell types and cell states, we performed additional benchmarking that used more complex datasets (**Figure 2E-F**). To increase the complexity of the classification task, we chose two datasets (Baron et al.^32^ and Tabula Muris mouse lung dataset^33^) that contain a larger number of cell types and overlapping phenotypes (cell states) that are difficult to discern. We included an additional 5 methods (Cell-ID^12^, AUCell^11^, Vision^34^, CellAssign^9^, Scmap-cluster, and CHETAH^35^) to compare the performance of scDECAF with a greater variety of reference-free tools, including probabilistic and rank-based methods, against reference-based methods for more diverse datasets.

In Baron Pancreas (**Figure 2E**), scDECAF was the third with the largest median F1 score after SingleR (first) and Cell Assign (second). While SingleR and CellAssign failed to annotate the rare Schwann cells, scDECAF recovered more than 75% of this population. In general, we see that most methods can correctly annotate some rare cell types of the pancreatic islet, such as stellate and delta cells, but not others, specifically epsilon. Many methods also fail to annotate mast, schwann, and endothelial cells, while scDECAF could correctly annotate these cell types. Among the methods considered, the per-cell-type F1 scores for scDECAF and Cell-ID had low variance and had a F1 of less than 0.2 for one cell population only - the rest of the methods failed to correctly annotate two or more cell types.

In the mouse lung dataset from the Tabular Muris consortium (**Figure 2F**), all methods but Garnett had a median F1 larger than 0.8, annotating a reasonably large proportion of cells accurately. We observed that reference-based methods generally perform better in distinguishing difficult-to-discern cell states such as classical versus non-classical cell states compared to methods relying on marker lists for cell annotation. None of the methods could recover the mast cell population in the Tabula Muris mouse lung cell dataset, potentially due to high degree of similarity to monocyte cells. The reduced performance of scDECAF and other marker list-based methods may be explained by the current lack of appropriate marker gene sets for mast cells and monocytes, and different cell states (classical and non-classical) for monocytes. Additionally, to demonstrate applications of the method in spatial transcriptomics, we applied scDECAF to annotate cell types in a slide-tags single-nucleus RNA (snRNA) of the human prefrontal cortex^36^ (**Figure 2G**). scDECAF was able to annotate cell types in this spatial transcriptomic data with 92% accuracy (**Supplementary Figure 1**).

A notable observation is that the performance of some reference-free methods - such as scDECAF, Vision, AUCell and CellAssign - was generally comparable to reference-based methods that were designed for cell type annotation. Our benchmarking illustrates that automatic cell type annotation may not require entire reference datasets. This may be an advantage in some biological contexts where appropriate reference datasets are not available, or where it is a goal to look for unexpected cell types, for example, the presence of neural cells in pancreatic islets is possible due to Pancreas-Brain crosstalk^37^, but they are not pancreatic cell types and will be absent in pancreatic references. The flexibility of scDECAF allows the user to define new labels in testing hypotheses, discover context-specific phenotypes and investigate spatial patterns of gene signatures. This may involve the application of scDECAF in a cell type annotation workflow alongside reference-based methods, or as is discussed below, in the characterisation of continuous phenotypes.

### scDECAF captures transcriptomic signals during EMT

Phenotypic heterogeneity within cell types is observed in biology but is not captured by cell type annotation methods. scDECAF allows researchers to associate the phenotypic heterogeneity observed in their data with functional information contained in knowledge bases and gene signatures, independent of clustering annotations. To assess the ability of scDECAF scores to identify cells along a spectrum of continuous biological states we used a well-studied, constrained biological problem: epithelial-to-mesenchymal transition (EMT). We used time course data^38^ (**Supplementary Figure 2**) where the authors exposed cancer cell lines to EMT inducers and performed scRNA-sequencing across 5 timepoints during EMT (0, 8 hours, 1 day, 3 days and 7 days) as well as 3 additional timepoints following the withdrawal of the EMT inducer (8 hours removed, 1 day removed, 3 days removed). We would therefore expect cells to lie on a continuum of EMT phenotypes and for scDECAF scores to reflect this transition. We used a pseudobulk differential expression analysis to define enriched gene sets that could be used to demonstrate the ability of scDECAF scores to reflect phenotypic heterogeneity.

A DE analysis between pseudo-bulked epithelial cells (‘Control’ cells at time point 0) and pseudo-bulked mesenchymal cells (‘EMT’ at days 3 and 7) revealed 800 differentially expressed genes (**Supplementary Table 1)**. The top 50 up- and down- regulated genes (**Figure 3A**) show a clear separation of control and EMT time points (highlighted). There is some variation in expression between other timepoints, showing the continuous nature of the EMT process, and variation between cell lines demonstrating the context specificity of EMT induction^38^. To minimize this variation and allow a clearer interpretation of enriched gene sets, we continued to restrict our comparison to Control and EMT timepoints (0 and 3-7 days). The 588 enriched gene sets (**Supplementary Table 2**) from this analysis reflect the expected phenotypes of cells undergoing TGFB1-induced EMT. Enriched gene sets include those that capture downstream TGFB1 signaling and process of EMT in cancers (**Figure 3B**). These expected transcriptomic shifts are observed in scDECAF phenotypes scores (**Figure 3C**), where score separations are consistent across cell lines (**Supplementary Figure 3**).

**Figure 3:**
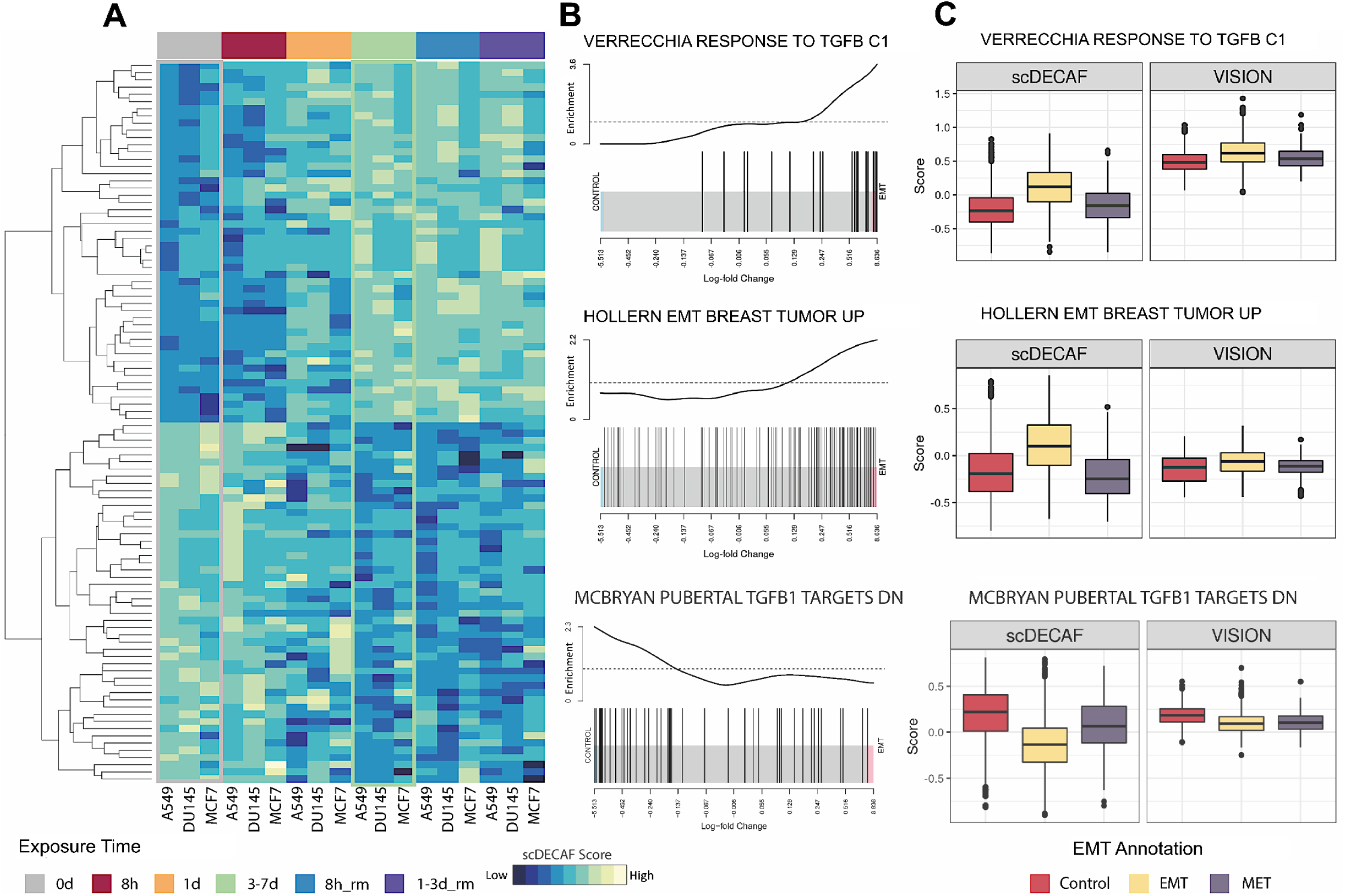
Correspondence between single-cell and pseudo-bulk methods for gene set analysis. **A.** Heatmap of batch-corrected logCPM expression values for pseudo-bulked samples of three cell lines (A549, DU145, MCF7) across 6 time points (0d, 8h, 1d, 3-7d, 8h removed, 1-3d removed) of a TGFB1-induced EMT time course. Heatmap is subset to the top 50 upregulated and 50 downregulated genes identified in the differential expression analysis between Control (0 days) and EMT (3-7 days) groups. Pseudo-bulk samples are aggregates of each time point within each cell line and columns are ordered by time course. **B.** Gene set analysis barcode plots for three gene sets from the C2 collection of MSigDB. Genes are horizontally ranked by log-fold change from the differential expression analysis between Control and EMT groups (See Methods) and the trace lines above the bars indicate enrichment. **C.** Boxplots of scores calculated by scDECAF and Vision for the same three gene sets in B). Scores are for cells across all three cell lines (A549, DU145, MCF7) at the Control (0 days exposure to TGFB1), EMT (3 & 7 days exposure) and MET (3 days removed from TGFB1) time points.

To illustrate the importance of per-cell scores that capture phenotypic variation in a dataset, we compared the distribution of scDECAF scores with those produced by Vision^34^ (**Figure 3C**). Although Vision scores show some variation between phenotypes, scDECAF scores show a stronger, more distinct difference. The mesenchymal cell populations (3-7 days) that have undergone EMT have observable differences in score for the Verrecchia TGFB1 and EMT breast tumor gene sets, compared to the Control (0 days) group of cells and to a lesser extent the final time point in the experiment (MET, 1-3d removed). The inverse is observed for the Mcbryan signature of downregulated TGFB1 targets that was down-regulated during EMT. This demonstrates that meaningful variation of per-cell scores capture phenotypic heterogeneity and enable the exploration of relevant biology.

### Discovery of biologically relevant gene programs in an Ovarian Cancer dataset

Single-cell transcriptomic data can contain diverse cell types and phenotypic heterogeneity that is difficult to summarize using pairwise comparisons between groups of cells. Few tools provide gene set scores for individual cells, where these scores are metrics that represent either some statistical test or aggregated expression for each gene signature^11,34,39–42^. Further, existing methods don’t prioritize gene sets for the user in a way that summarizes the major sources of variation in the dataset. Here we demonstrate how scDECAF’s “discovery” (sparse gene set selection) mode, that is, the sparse gene program selection mode, identifies biologically relevant gene sets out of the thousands of gene sets in the C2 collection of MSigDB.

We analyzed an ovarian cancer dataset that contains malignant cells, fibroblasts, and immune populations from the ascites effusions of 6 patients^43^ (**Supplementary Figure 4**). scDECAF pruned 5,529 gene sets down to 91, and the gene set scores were further investigated using the cell type annotations, patient information and tSNE coordinates reported in the paper (**Figure 4A**). We found that many of the 91 prioritized gene sets showed elevated scores in expected cell types, such as malignant cells expressing ZEB1 targets (AIGNER_ZEB1_TARGETS) or stem cell transcriptional programs (YAMASHITA_LIVER_CANCER_STEM_CELL_UP), fibroblasts expressing VEGFA targets (WESTON_VEGFA_TARGETS_12HR), and B cell clusters having elevated scores for antigen presentation (PELLICCIOTTA_HDAC_IN_ANTIGEN_PRESENTATION_UP). There are also subtle differences observed across patient clusters, such as stronger expression of ZEB1 targets in Patient 3 malignant cells compared to other patients. Since ZEB1 suppresses its targets to downregulate epithelial genes and promote invasiveness, increased expression of ZEB1 targets would indicate maintenance of epithelial cell identity. We also observe that fibroblasts in Patient 5.1 that have relatively higher scores for WESTON_VEGFA_TARGETS_12HR, indicating stronger VEGFA signaling in these cells.

**Figure 4:**
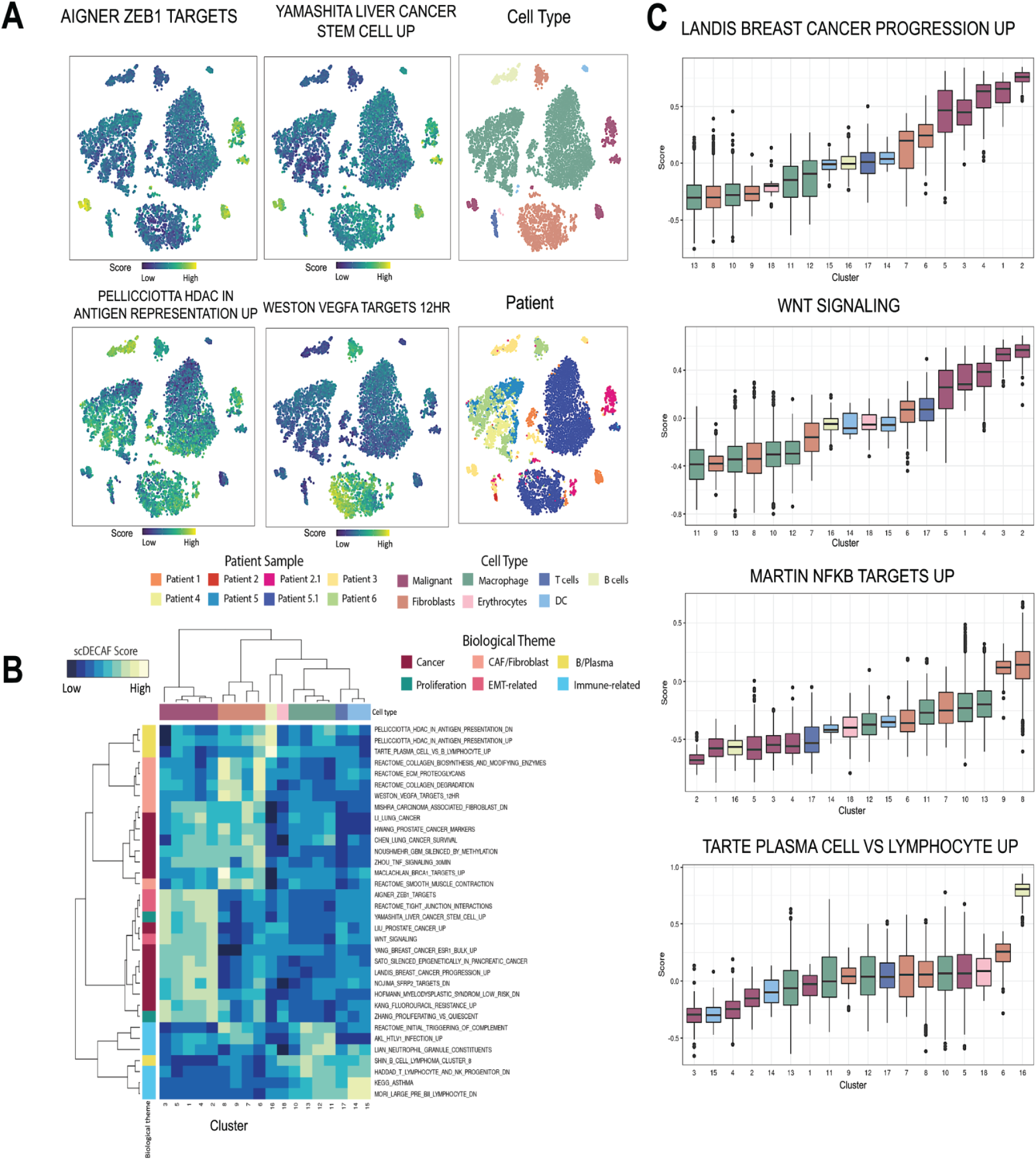
The scDECAF “discovery” pipeline reveals known and novel biology. The scDECAF “discovery” pipeline was applied to an ovarian cancer dataset, providing per-cell scores for 91 pruned gene sets. **A.** Using the tSNE embedding from the original paper^43^ we highlight the correspondence of scDECAF scores with patient and cell type clusters; tSNEs are coloured by cell type, patient, or scDECAF score. Cluster and cell type annotations from the original paper were used throughout the figure. Cell type colors are used in B and C. **B.** The heatmap shows the mean scDECAF score for each cluster for a selection of 34 gene sets, coloured by biological theme (rows) and cell type (columns), following hierarchical clustering. **C.** Boxplots show the scDECAF scores within each cluster for gene sets of interest, coloured by cell type.

For gene sets with recognisable themes, we summarized the scores with hierarchical clustering to show that scDECAF has selected biologically relevant gene sets that broadly correspond to the phenotypes of cell type clusters (**Figure 4B**). For example, cancer-related gene sets were associated with malignant cell clusters, while gene sets relating to ECM remodeling and cancer-associated fibroblast phenotypes clustered with fibroblast clusters.

We also investigated the data by cluster, using the cluster annotations provided by the authors, to show that scDECAF captures phenotypic heterogeneity between clusters of the same cell type (**Figure 4C; Supplementary Figure 5**). For example, while malignant cell clusters scored highly for cancer related gene sets (LANDIS_BREAST_CANCER_PROGRESSION_UP & WNT_SIGNALING) consistently across patients, scores were more elevated in some clusters. Clusters 1, 4 and particularly 2 have higher LANDIS_BREAST_CANCER_PROGRESSION_UP scores and shorter box plots that indicate a tighter score distribution within the cluster. This may reflect an increased aggressiveness in the cancers of these patients. Similarly, two cancer clusters (2 and 3) with relatively higher WNT_SIGNALING scores might reflect patient-specific differences in Wnt signaling. Some cell types displayed even stronger score differences between clusters, such as MARTIN_NFKB_TARGETS_UP scores in fibroblasts, where clusters 8 and 9 are the only clusters with score distributions above zero and clusters 6 and 7 are much lower for this pathway. This phenotypic heterogeneity across fibroblast populations from different patients may reflect different levels of NFKB signaling across tumor microenvironments. In the case of TARTE_PLASMA_CELL_VS_B_LYMPHYOCYTE_UP scores, which are strongly elevated in the B cell cluster, gene set scores may help to confirm cell type annotations or assist in distinguishing between phenotypically similar cell types, such as plasma and B cells, where annotation may have failed.

### Sparse gene program selection coupled with reference mapping reveals multicellular processes associated with severe COVID-19

Recent computational developments^44–46^ have empowered understanding the biological processes in disease progression at single cell resolution through construction of healthy single cell references and mapping query disease samples onto them for contextualization. Recent findings suggest that disease-specific cell states in case-control scRNAseq datasets can be optimally found by mapping to a healthy reference atlas^47^ . Here, we construct a CITE-seq reference of healthy PBMC cells by totalVI^48^ and map a CITE-seq PBMC dataset containing cells from healthy individuals and COVID-19 patients^49^ (**Figure 5A**) to identify cell types in the query. Differential abundance analysis using Milo^50^ identified significant compositional changes in Platelets, Plasmablasts, pDCs, Erythrocytes and cell states in CD14 Monocytes that were increased in severe COVID-19 compared to healthy controls (**Figure 5B, Supplementary Figure 5 A-B**).

**Figure 5:**
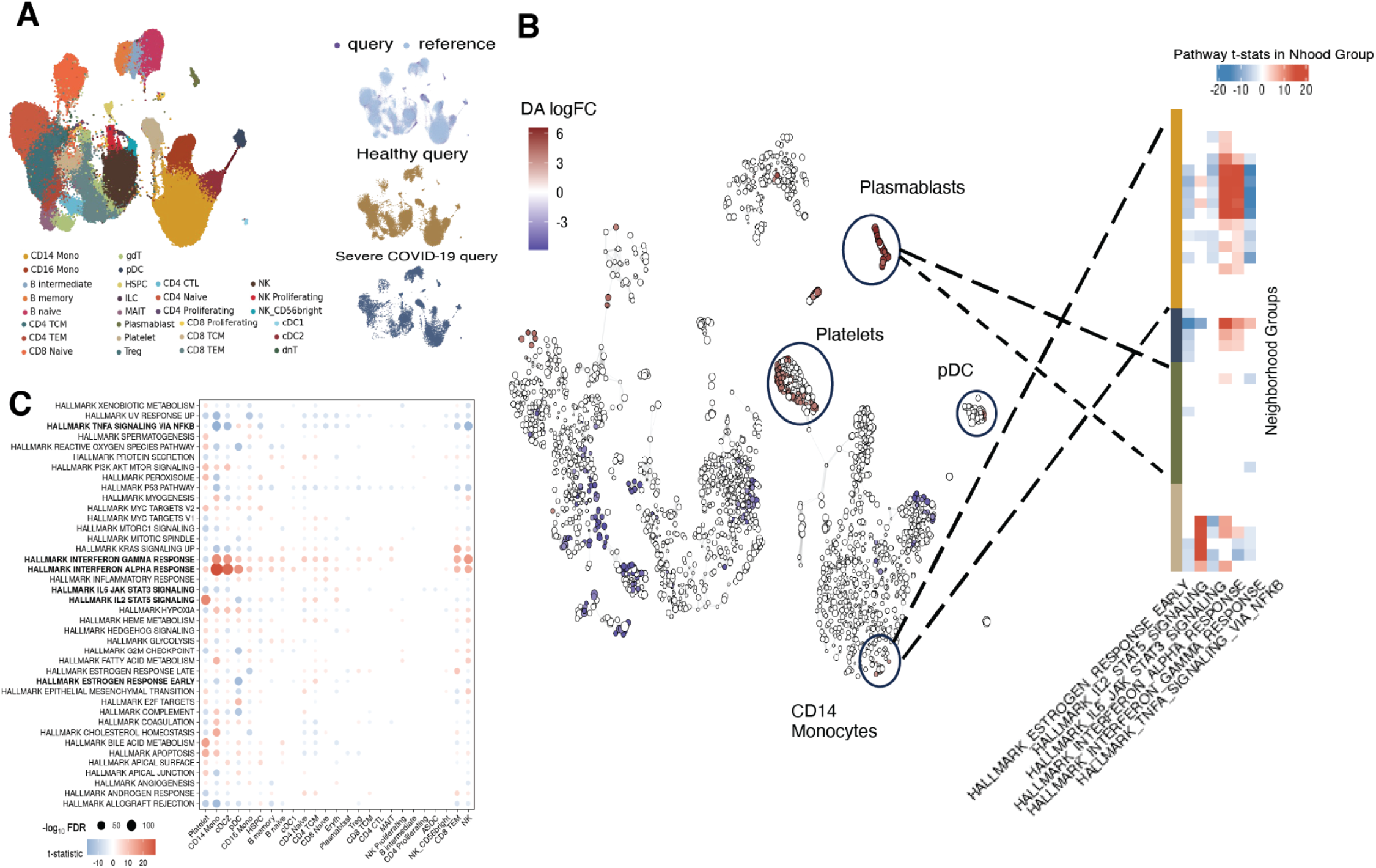
Application of scDECAF to PBMC cells from COVID-19 patients and healthy individuals mapped onto a healthy CITE-seq PBMC reference delineates compositional and cell-intrinsic gene program changes associated with disease severity that are coordinated across cell types. **A.** UMAP representation of an integrated healthy PBMC reference and a COVID-19 case-control CITE-seq dataset coloured by cell type, query and disease status. **B.** Milo Neighbourhood Graph constructed from the UMAP in A subset to cells from the COVID-19 case-control query, highlighting differentially abundance cell states in severe COVID-19 compared to healthy individuals and changes in transcriptional activity in a number of Hallmark gene programs selected by scDECAF within those cell states. CD14 Monocytes, Platelets and pDCs undergo both compositional and transcriptional shifts in COVID-19, while Plasmablasts only undergo compositional changes in the query dataset. **C.** Heatmap of t-statistics for differential pathway activity in COVID-19 in cell neighborhoods identified in B for Hallmark gene programs selected by scDECAF. Pathway activity is quantified by scDECAF. For each cell type and pathway, the most significant t-statistic over all cell neighborhoods was chosen. Color represents the sign of the t-statistic, size represents the significance of the test (-log_10_ FDR).

We applied scDECAF to the embedding of query cells post reference mapping to identify biologically relevant gene programs from the MSigDB Hallmark collection and score the transcriptional activity of these gene sets. We used the Milo graph to obtain neighborhood groups by merging statistically significant neighborhoods that had consistent log fold-changes and summarized the activity levels per neighborhood group. We then tested for differential activity of the selected gene sets to identify gene programs associated with severe COVID-19 within each neighborhood group (**Supplementary Figure 5C**). The differential activity analysis on the K-nearest neighbor Milo graph revealed that while CD14 Monocytes, Platelets and pDCs had undergone both compositional and gene program changes in severe COVID-19, Plasmablasts had only undergone a composition change (**Figure 5B**).

The prioritized Hallmark gene programs identified by scDECAF - that were also found to have differential activity in severe COVID-19 at the cell state level (**Figure 5C**) - were consistent with previous reports^49,51–53^ on deregulation of IL6-Jak-STAT3 signaling, inflammatory response, Interferon alpha response, interferon gamma response, IL2-STAT signaling, TNFA signaling via NFKB, heme metabolism, and G2M checkpoint in COVID-19 in pDCs and CD14 Monocytes (**Figure 5C**). In addition to observing coordinated activity of interferon alpha and gamma response, we also detected coordinated activity of TNFA signaling via NFKB (**Figure 5C**) in cell types that had differential activity of interferon alpha and gamma response in severe COVID-19 patients compared to healthy individuals, suggesting a multicellular program associated with severe COVID-19. For platelets specifically, we found that IL2-STAT5 signaling had differential activity in cells from patients with severe COVID-19. The role of STAT5 in platelet production is already established in hematopotesis^54^, indicating the relevance of the IL2-STAT5 program in platelets from COVID-19 patients identified by scDECAF. Since most patient samples were 9-15 days post infection, detection of reduced differential activity of Early Estrogen Response in pDC and CD14 Monocytes was also a reasonable finding given its role in repression of interferon signaling^55^ . This suggests that the impaired Early Estrogen Response in these cell types interfered with modulation of immune response and led to the development of severe disease stage.

These findings collectively suggest that scDECAF can be used for post-hoc identification of cell type-specific gene programs and multicellular processes, that is, coordinated gene programs across cell types, that are associated with disease or perturbations. When applied downstream of reference mapping and differential abundance workflows, scDECAF can be used to decouple cell-intrinsic gene program activity from cell compositional changes.

## Discussion

In this work, we demonstrated that scDECAF can identify cell types and states, efficiently query gene programs and aid biological interpretation using gene set representations.

We applied scDECAF to cell type annotation benchmark datasets from four tissues, using tissue-specific cell type markers from databases such as CellMarker. Where good quality marker lists are available, the accuracy of scDECAF in cell type annotation was comparable to annotation transfer from a reference. In general, we observed that reference-based cell type annotation algorithms outperform marker list-based methods, largely due to some cell types being difficult to distinguish and lacking a high quality marker list. However, marker list-based methods can empower the discovery of cell types and states in biological contexts where they have not been previously found or reported, allowing for the identification of cell types not present in reference datasets. We extended our evaluations to spatial transcriptomics where we were able to annotate cell types with high accuracy where spatial gene signatures were available.

We used molecular signatures from MSigDB to infer gene program expression and demonstrate that scDECAF scores could distinguish subpopulations between EMT time points for known differentially regulated gene sets. Additionally, we demonstrated that scDECAF can recover perturbed gene programs in PBMC cells from Lupus treated with IFN-beta. scDECAF also identified B cell-specific programs in IFN-beta-treated cells that could not be found by conventional GSEA analysis. Similar to other score-based methods, scDECAF is cell type-agnostic; it can reveal gene programs that are present across cell types and those that exhibit cell type-specific activity. Therefore, scDECAF should not be used to replace GSEA where cell type-specific enrichment is of interest.

In our experiments, we observed that the learned vector representations of cellular gene expression profiles and gene sets strongly depend on the number of input gene sets (**Supplementary Note 1**). This implies that the strength (magnitude) of the scores can change as gene sets are added or removed from the input list. To mitigate this issue, scDECAF uses a cell embedding to select the subset of programs that are most relevant for reconstruction of the embedding. To show how this works in practice we applied scDECAF to an ovarian cancer dataset using thousands of signatures from MSigDB to identify 91 biologically relevant programs. In this example, we demonstrated how scDECAF can be used to discover patient-specific phenotypes and aid the interpretation of a heterogeneous cancer dataset.

Rood et al^56^ highlighted a need for algorithms for efficient interrogation of gene programs, cell types and state among the key computational challenges for cell atlases in medicine. In addition to limited scalability to large-scale atlases, ranked- and factorisation-based gene program scoring methods would work optimally with batch-corrected data which may not be available for integrated atlases. While interpretable deep generative models are scalable to cell atlases, they can not be applied to existing integrated atlases or reference mapping. Application of scDECAF to PBMC cells from COVID-19 patients and healthy individuals mapped onto an existing healthy CITE-seq PBMC reference, in conjunction with differential gene program activity analysis, delineated compositional and cell-intrinsic gene program changes associated with disease severe COVID-19 that are coordinated across cell types.

The sparse gene program selection module of scDECAF only requires a cell embedding, which is available for integrated atlases or existing references, making our method versatile to a range of analyses. Additionally, scDECAF can be applied to cell-specific Bayes Factors from deep generative models or cell-specific scores from matrix factorization-based methods. This work can, therefore, complement reference mapping and differential abundance analysis frameworks, and other conventional gene programs activity inferring tools. We, therefore, expect that scDECAF can be used for efficient query of cell types, states and programs in single cell RNA sequencing data, particularly in cell atlases.

## Materials and Methods

### scDECAF Learns vector representations of cells and gene sets by Canonical Correlation Analysis (CCA)

Canonical Correlation Analysis^18^ involves finding a lower-dimensional representation of two datasets *X* and *Y*, where the correlation between *X* and *Y* is maximized. scDECAF uses CCA to find lower-dimensional representation of cells (*X*) and a binary matrix encoding gene sets (*Y*) in scRNA-seq data, where gene expression in the cells is maximally correlated with the patterns induced by genes in the gene sets. Let *n* represent the number of genes, *p* number of cells in *X* and *q* the number of gene sets in *Y*. Assume that the columns of *X*_*n*×*p*_ and *Y*_*n*×*q*_ have been centered and scaled. CCA finds *u*_*p*×*r*_ and *v*_*q*×*r*_, *r* < *min*(*p*, *q*), that maximize *Cor*(*Xu*, *Yv*) - that is, it solves

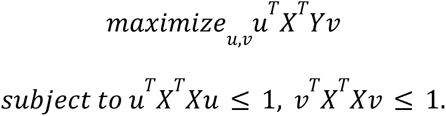

The vectors *u* and *v* are lower-dimensional representations of gene expression data and gene set binary encodings respectively, and are called the canonical variates. We use Witten et al.^19^ (2009), implementation of CCA to find *u* and *v*. These vectors are then *L*_2_ normalized:

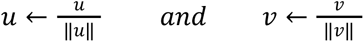

#### Cell-gene set assignment by similarity of vector representations in the vector space

scDECAF operates in two modes: cell type annotation and phenotype scoring, with the additional option of performing gene set selection (sparsity inducing mode) with phenotype scoring. In phenotype scoring mode, a score is assigned to each query gene set per cell. This score is determined by the similarity of vector representation of cells and gene set in the latent space learned by CCA. For example, the score *s_cj_* for cell *c* and gene set *j* is determined as

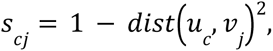

where *dist*(*u*, *v*) is the Euclidean distance, and 1 ≤ *c* ≤ *p*, 1 ≤ *j* ≤ *q*. Note that since *u* and *v* are *L*_2_ normalized, *s_cj_* is equivalent to cosine similarity between vector representation *u_c_* of cell *c* and the vector representation *v_j_* of gene set *j* on the latent space. Therefore, *s_cj_* measures the strength of association between the label of the *j*th gene set and gene expression in cell *c*.

In the cell type annotation mode, the label of the closest gene set *v_d_*

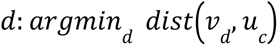

is assigned to cell *c*.

The dimensions of the latent space learned by CCA only capture cell-gene set similarities and cell-to-cell transcriptional similarities are lost in this latent space. Dimension Reduction techniques such as tSNE and UMAP produce embedding of the cells that preserve local and global structures in scRNA-seq data. We, therefore, refine the initial cell-gene set assignments, that are determined based on the distance in CCA space, according to a local or global neighborhood of the cell on a gene expression embedding, say the UMAP embedding of the cells. This is motivated by the fact that cells with similar transcriptional profiles that are close to each other on UMAP embedding should be more likely to have identical labels.

Label refinement for cell *c* is performed by replacing the initial label with the label supported by the majority of cells in a small or large neighborhood of the cell, *N*_c_, weighted by the distance of the cells from cell *c*. For each query cell c, we extract the *k* nearest neighbors *N_c_*. We compute the standard deviation of the nearest distances:

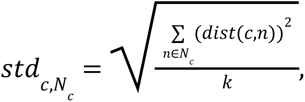

where *dist*(*c*, *n*) is the Euclidean distance of the query cell *c* and its neighbors *n* on the (e.g. UMAP) embedding. We the apply the Gaussian Kernel to distances as follows:

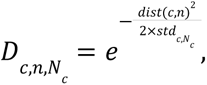

The probability of assigning each label *y* observed in the neighborhood to the query cell *c* is computed as:

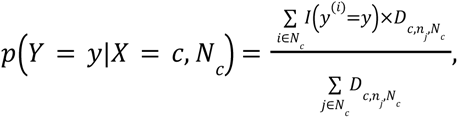

where *y*^(*i*)^ is the label of i-th nearest neighbor. The label with maximum probability is considered as the refined label. that is:

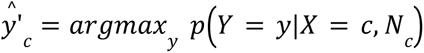

The uncertainty in the assigned label is estimated using:

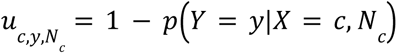

finally, if uncertainty is larger than a user-specific value, κ, that is cells in *N*_*c*_ are highly heterogeneous in labels, the cell label is assigned as “Unknown” - that is:

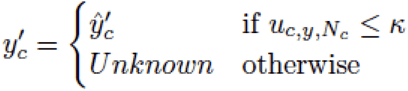

### A model-based strategy for selection of relevant gene sets

The learned CCA space is dependent on the number of gene sets included in the list of query gene sets. The cell-gene set assignments, distribution of scores and the strength of associations will vary as certain gene sets are included in, or excluded from the query list. For optimal performance, the gene sets most relevant to the data should be selected from a query list during sparse gene set selection mode.

We define the gene set selection problem as a multi-response penalized linear regression problem, where mean expression of genes in each gene set is included as a predictor. The coordinates of the cells on each dimension of the UMAP embedding are the response variables (hence why the model is a multi-response regression: response 1, say *z*_1_, is the coordinate of the cells on the first dimension of the UMAP, and *z*_2_ is the coordinate of the cells on the second dimension of UMAP). We use Lasso and Elastic-Net Penalized Generalized Linear models (GLM) of Friedman et al.^20^(2010), which solves the following problem:

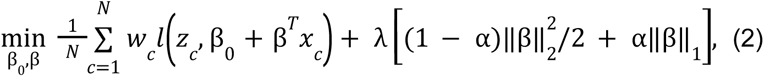

where *z*_c_ is the coordinate of cell *c* on a UMAP dimension (i.e the response), *x*_c_ is the scores for gene sets in cell *c* where score is simply the average expression of genes in the gene set, and λ is the penalty (regularization) term. The optimal penalty, λ, is determined by 10-fold cross validation, whereas α is specified by the user and determines if a Lasso (α = 1) or Elastic-Net (α = 0. 5) model should be fitted . In practice, the model in (2) selects a subset of gene sets that minimizes the mean squared error of the predicted UMAP coordinates from the observed UMAP coordinates - that is, gene sets that are predictive of the position of the cell on UMAP embedding. This results in selection of gene sets that are expressed or inactivated uniformly in all cells in the cell clusters. These gene sets, therefore, are likely to represent transcriptional programs governing a cluster of cells. The model in (2) is fitted simultaneously to all response variables when fitting a penalized GLM model with the assumption that each UMAP dimension follows a normal distribution.

### Datasets

We report an extensive list of datasets at Appendix Table S1.

### Gene set analysis in interferon stimulated Lupus PBMC

Single cell RNA counts from control and IFNβ-stimulated lupus PBMC cells^21^ were normalized by SCTransform^57^ and 2000 HVG. We then ran scDECAF with Hallmark gene sets^22,58^. The optimal shrinkage value for sparse gene program selection was determined by inspecting the reconstruction (of the UMAP embedding) error plot. We set the sparsity regularizer parameter lambda to exp(-4), which corresponded to the lowest reconstruction error. Once the set of gene programs resulting in optimal reconstruction were identified, we ran scDECAF on the selected set to compute gene program activity scores per cell. To compare the results with standard gene set enrichment analysis, we used fgsea^59^ to identify Hallmark gene sets enriched in B cells in response to interferon stimulation.

### Cell type labeling benchmarks

For the cell type annotation benchmarking we downloaded and processed 8 datasets, from 4 tissues (Table 1 Appendix). These datasets were downloaded and processed using Seurat, including highly variable gene (HVG) selection and SCTransform normalization, where the corresponding reference dataset was used when required by a method. Datasets were subset to an appropriate number of HVGs (See Description of datasets and processing in Supplementary information). Since reference and query datasets did not always contain the same cell types, labels were either mapped if there was an appropriate matching cell type (**Supplementary Tables 3-6**), or excluded.

In our evaluations, we found that scDECAF works well with 2000 HVGs (this number is also recommended for data integration). However, if there is a strong similarity between gene sets, e.g. cell types or phenotypes are indistinguishable, or there is a minimal overlap between genes in the gene sets and the selected HVG genes (that is, most of the genes in the gene sets are not highly variable), users may select a larger number of HVG for scDECAF analysis depending on the complexity of the dataset. In practice, we observe that cell clusters are more discernible with 2000 HVG genes. We used UMAP as the input embedding for all cell type annotation benchmarking.

In cell type annotation mode, the initial labels of the cells undergo a refinement, or smoothing, based on a neighborhood *N_c_* around the cell. This neighborhood is determined by the value of *k*: if *k* is small, labels are refined considering local similarity of cells. If *k* is large, the global similarity of cells is considered for label refinement.

We evaluate the accuracy of cell type classification algorithms by comparing the predictions to previously published cell type labels. Accuracy is determined by F1-score and label mapping scheme described in (Supplementary Tables 3-6) :

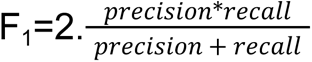

### Algorithms for benchmarking

#### Garnett

Garnett^10^ finds some seed cells for each cell type based on a list of input marker list. It then trains a classifier on the data using the seed cells. The classifier is then used to predict the identity of the cells. We used the marker lists provided on their github page https://cole-trapnell-lab.github.io/garnett/classifiers/

#### scmap

Scmap^8^ is a reference-based cell type annotation technique that computes the similarity of cells in the query dataset with the cells in the reference by cosine similarity. The cells in the query dataset are assigned with identical labels as cells if the reference, if similarity is larger than a threshold. Label assignment can be performed at cell cluster or individual cells level. We used the Biocondutor implementation of scmap https://www.bioconductor.org/packages/release/bioc/html/scmap.html

#### SingleR

SingleR^7^ is also a reference-based cell type annotation algorithm. SingleR annotates cells in the query dataset based on the Spearman correlation of gene expression with the expression levels in the reference dataset. We used the Biocondutor implementation of SingleR https://www.bioconductor.org/packages/release/bioc/html/SingleR.html

#### Seurat v4

We followed Seurat’s official vignette for mapping and annotation of query datasets https://satijalab.org/seurat/articles/integration_mapping.html

#### cellAssign

cellAssign^9^ probabilistically assigns each cell in a single-cell RNA-seq dataset to a cell type based on a priori known markers for cell types. We used the github implementation here https://github.com/Irrationone/cellassign

#### AUCell

AUCell^11^ allows the identification of cells with active gene sets (e.g. signatures, gene modules…) in single-cell RNA-seq data. AUCell uses the Area Under the Curve (AUC) to calculate whether a critical subset of the input gene set is enriched within the expressed genes for each cell. We used the Bioconductor implementation of AUCell. https://www.bioconductor.org/packages/release/bioc/html/AUCell.html

#### CelliD

CelliD^12^ is a tool for unbiased extraction of Single Cell gene signatures using Multiple Correspondence Analysis. We used the Bioconductor implementation of CelliD. https://www.bioconductor.org/packages/release/bioc/html/CelliD.html

#### Vision

Vision^34^ is a gene set scoring method that provides a representative score for each gene signature in every cell using normalized aggregate expression. This software is available on github: https://github.com/YosefLab/VISION

#### CHETAH

CHETAH^35^ is a scRNA-seq classifier. CHETAH creates a classification tree by hierarchical clustering of the reference data. CHETAH classifies the input cells to the cell types of the references. We used the Biocondutor implementation here https://www.bioconductor.org/packages/release/bioc/html/CHETAH.html

### Phenotype scoring in the Ovarian Cancer and time-course EMT datasets

Datasets were downloaded from the NCBI Gene Expression Omnibus (GEO) database including 9,609 high-grade serous ovarian cancer cells from malignant ascites samples^43^ (GSE146026) and 8,419 cells from 3 cell lines (A549, DU145, MCF7) across 8 time points of EMT induction^38^ (GSE147405). From the time course data, we used only the TGFB1-induced samples and excluded the OCVA420 cell line due to low cell numbers across time points.

Non-expressed genes and duplicated gene names were filtered from both datasets and 3000 features (genes) were selected prior to analysis The normalized counts for the ovarian cancer dataset were otherwise used as provided by Izar et al.^43^, whereas Seurat was used to normalize the EMT time course dataset.

scDECAF was applied to both datasets to obtain phenotype scores across C2 gene sets from MSigDB^58,63^. When running scDECAF, 10 components were calculated, UMAP was used for input embeddings (**Supplementary Figures 2 & 4**) and standardization and k=10 were used. For the ovarian cancer dataset, scDECAF pipeline was run on discovery mode (sparse gene set selection) and gene sets were pruned down to 91 gene sets. To prune gene sets we used a lasso penalty *λ* = e^-2^ and UMAP as the input embedding.

Vision^34^ was used to obtain gene set scores for comparison with scDECAF scores. Counts were normalized using scater^60^ but not log-transformed, pooling within Vision was set to false and gene set scores were obtained for the same C2 gene sets used in the scDECAF analysis above.

### Pseudo-bulk differential expression analysis

Pseudo-bulk samples were obtained by aggregating each cell line at each time point to reflect a bulk RNA-seq experiment, giving 18 pseudo-bulk samples. Some time points were merged (3 & 7 days, 1 & 3 days removed) to balance pseudo-bulk sample library sizes. Pseudo-bulked samples were filtered and normalized using standard edgeR^61^ pipelines. Genes were retained only if they were expressed at a logCPM >= 0.5 in at least 10% of samples. Cell line batch effects were estimated using limma^62^ and removed for visualization. A quasi-likelihood framework was applied to identify genes that were differentially expressed between what we define as ‘Control’ and ‘EMT’ cells, corresponding to day 0 cells and day 3-7 cells, respectively. We used P-value and FDR thresholds of 0.05 and performed Benjamani-Hochberg P-value correction.

Competitive gene set enrichment analyses were also performed on pseudo-bulk samples using limma. Enrichment was performed with gene sets in the C2 collection of MSigDB, for gene sets with more than 10 genes.

### COVID-19 CITE-seq PBMC

Reference building and query mapping of the healthy CITE-seq PBMC and healthy and COVID-19 PBMC datasets were performed by totalVI^48^ as described in scvi-tools tutorial https://docs.scvi-tools.org/en/stable/tutorials/notebooks/totalVI_reference_mapping.html

The RNA modality of the query CITE-seq dataset was normalized by SCTransform and 2000 HVG. We then ran scDECAF with Hallmark gene sets. The optimal shrinkage value for sparse gene program selection was determined by inspecting the reconstruction (of the joint totalVI embedding) error plot. We set the sparsity regularizer parameter lambda to exp(-4.5), which corresponded to the lowest reconstruction error. This resulted in selection of 40 Hallmark gene sets. Once the set of gene programs resulting in optimal reconstruction were identified, we ran scDECAF on the selected set to compute gene program activity scores per cell.

We used Milo^50^ to carry out differential abundance analysis between severe COVID-19 and healthy query cells. Milo neighborhood graph was constructed using the 2-dimensional UMAP representation of the query data in the integrated embedding, setting the k-nearest neighbor parameter to 30. We used default parameters for all other remaining arguments. Differential abundance (DA) was determined based on Spatial FDR < 0.05. Neighborhoods with consistent DA logFC and spatial FDR < 0.05 were grouped using the groupNhoods() function. We then tested for the differential activity of the 40 selected gene sets in severe COVID-19 versus healthy in each neighborhood group using the testDiffExp() function in miloR, which calls limma under the hood akin to differential gene expression. These analyses were performed within query data, on the integrated embedding, that is post reference mapping, which is found to result in optimal detection of disease-specific cell states using healthy cell atlases^47^ . To summarize differential pathway activity over all the neighborhood groups in a cell type, we reported the most significant t-statistic (smallest adjusted P-Value from limma’s moderated t-statistic) in neighborhood groups of a cell type.

## Acknowledgements

We acknowledge the reviewers from the ICML 2021 Workshop on Computational Biology who provided very insightful feedback about an earlier version of this manuscript. We thank Yi Xie and Jack Finlay for helpful discussions.

## Author contributions

S.H conceived the project under supervision of M.J.D. S.H and H.J.W performed the data analysis with contributions from M.K, F.C and D.B. S.H wrote the code. S.H, H.J.W and M.J.D have written the manuscript. All authors have read and approved the manuscript.

## Data availability

All data used in this study is published and is cited throughout the text. We report an extensive list of datasets and accession numbers in Supplementary Information.

## Code availability

scDECAF is available as an R package at https://github.com/DavisLaboratory/scDECAF. R Markdown files to reproduce the results and figures are available at https://github.com/DavisLaboratory/scDECAF-reproducibility.

## Supplementary Materials

**Supplementary Figure 1.**
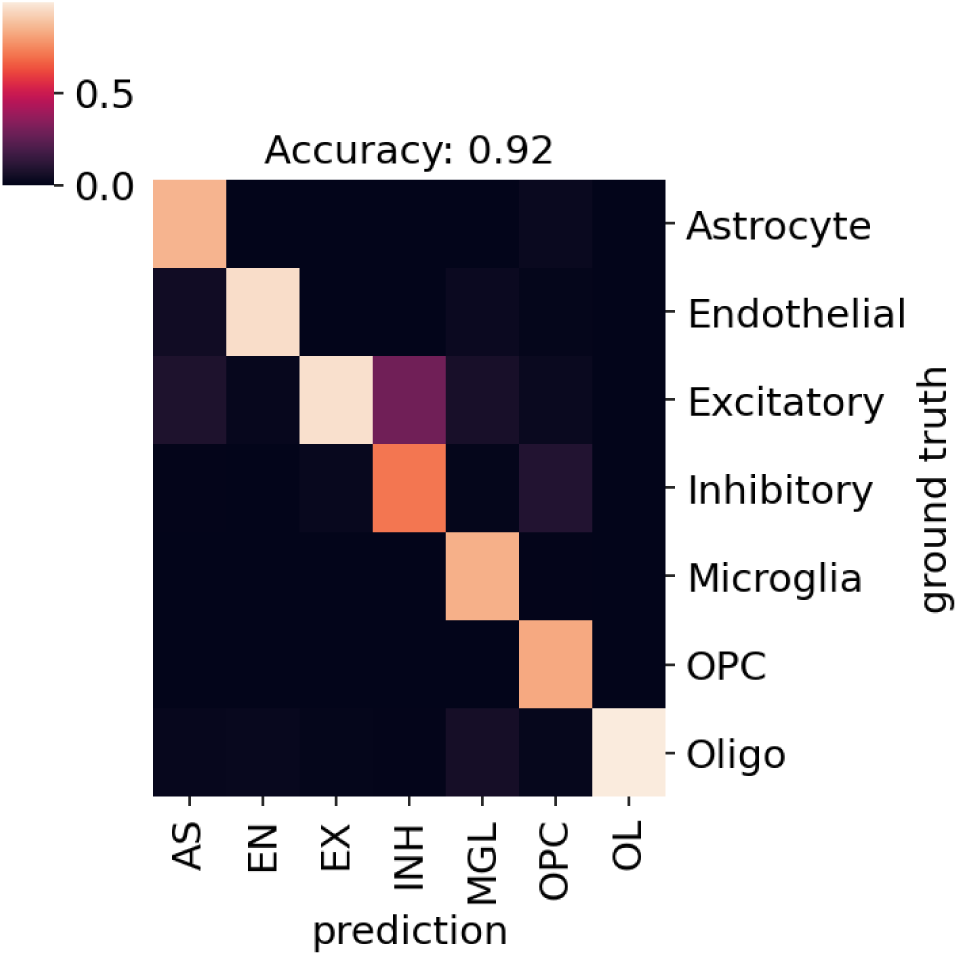
Confusion matrix for cell type predictions in the slide-tags snRNA from human prefrontal cortex

**Supplementary Figure 2.**
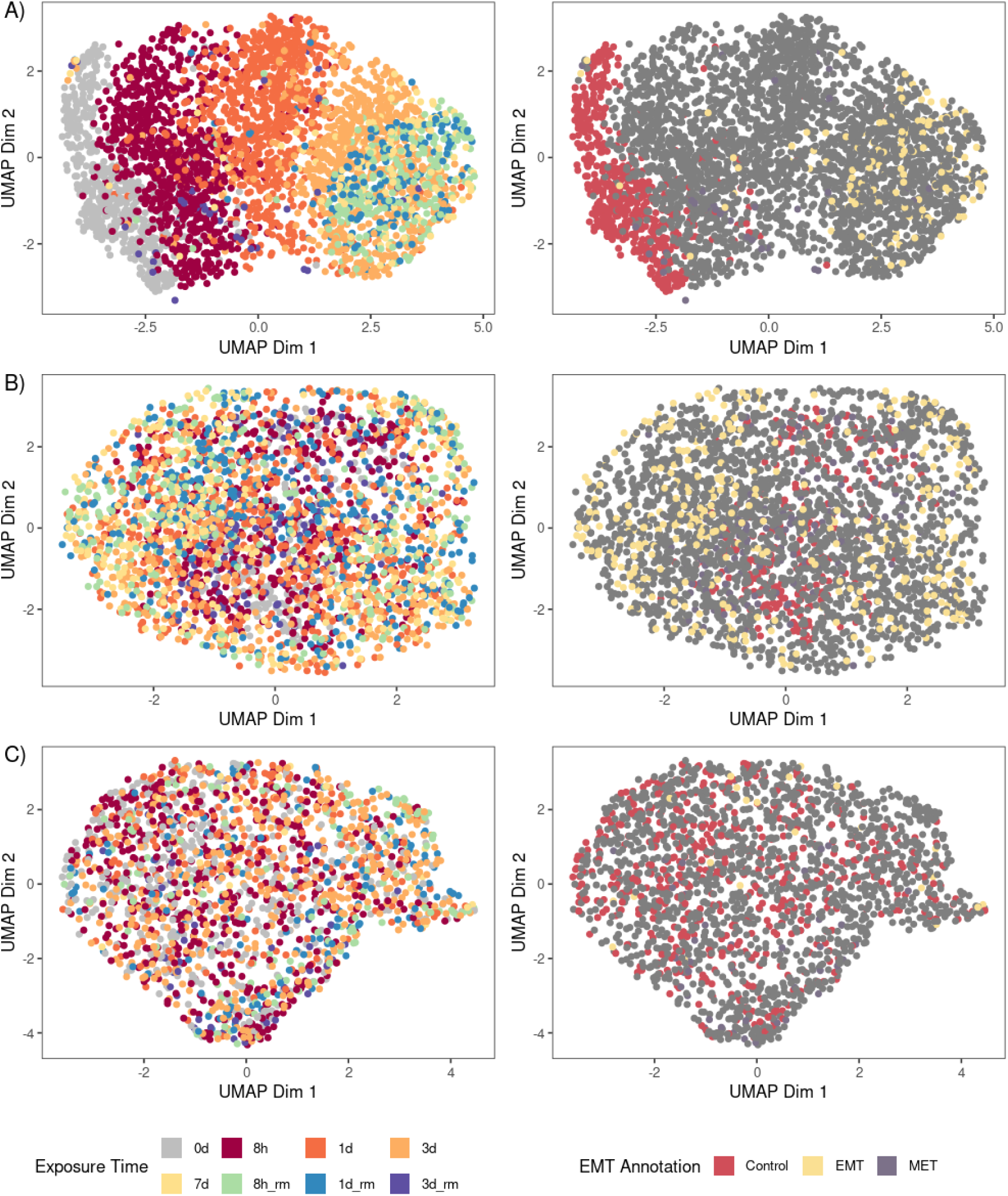
The UMAP embeddings used by scDECAF to calculate scores for each cell line - A549 **(A)**, DU145 **(B)**, and MCF7**(B)** - in the EMT dataset, using 3000 HVGs each. Cells are coloured by TGFB1 exposure time (left UMAP) and EMT status annotation (right UMAP) used for differential expression analysis.

**Supplementary Figure 3.**
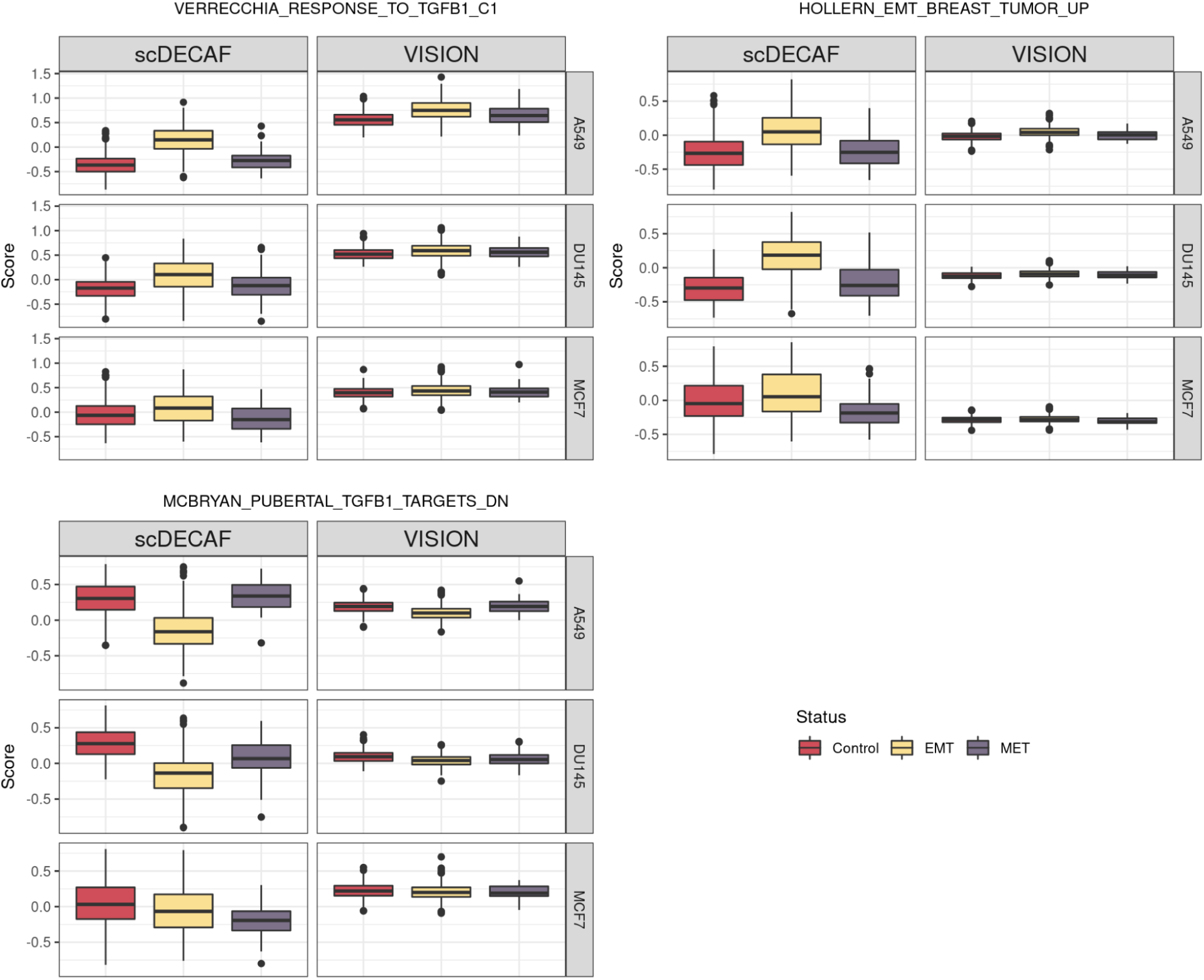
Boxplots of scores calculated by scDECAF and Vision for the same three gene sets in Figure 3, faceted by cell line. Scores are for cells across all three cell lines (A549, DU145, MCF7) at the Control (0 days exposure to TGFB1), EMT (3 & 7 days exposure) and MET (3 days removed from TGFB1) time points.

**Supplementary Figure 4.**
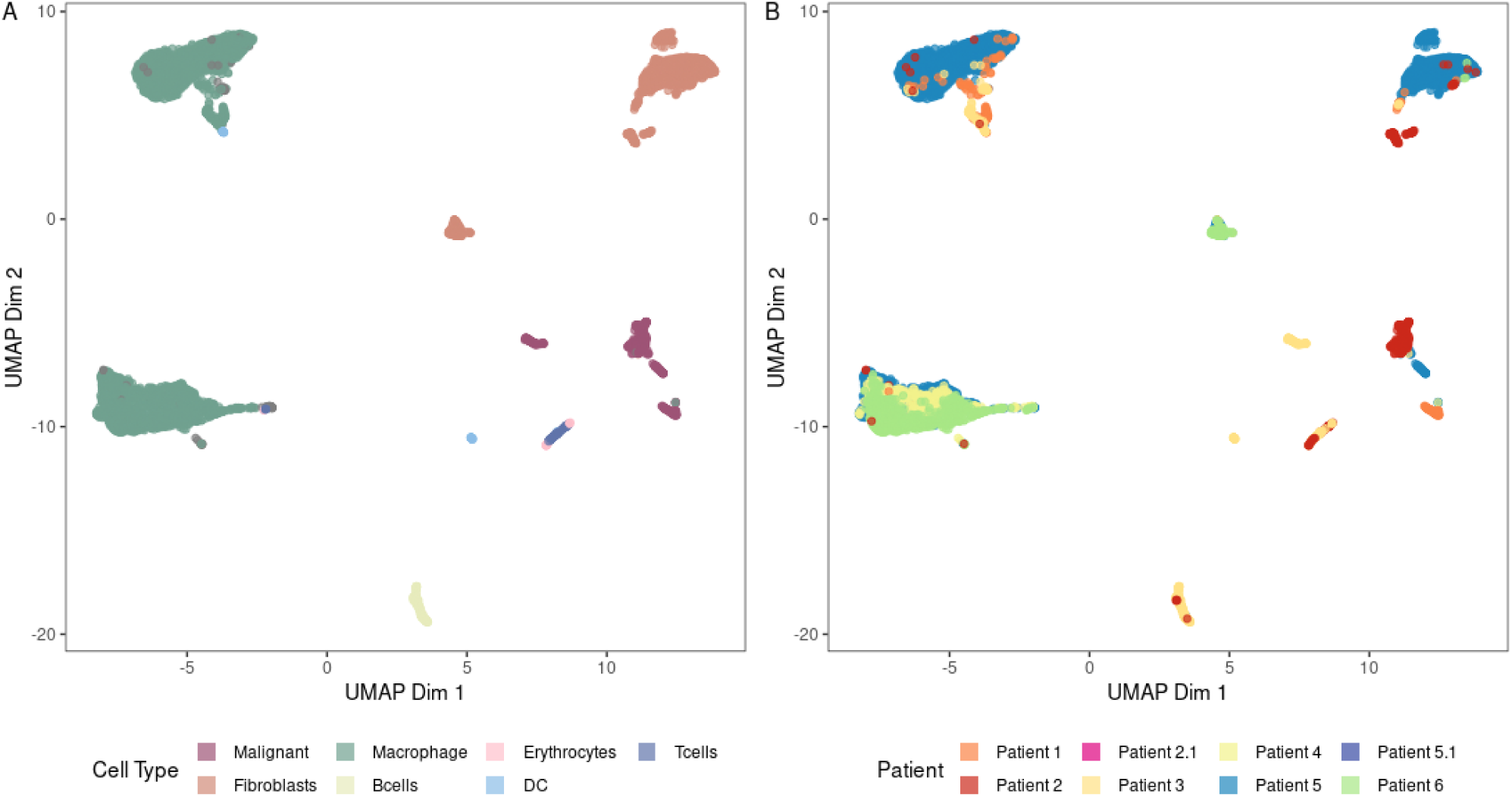
The UMAP embedding used by scDECAF to calculate scores for the ovarian cancer dataset, coloured by cell type **(A)** or patient **(B)**.

**Supplementary Figure 5.**
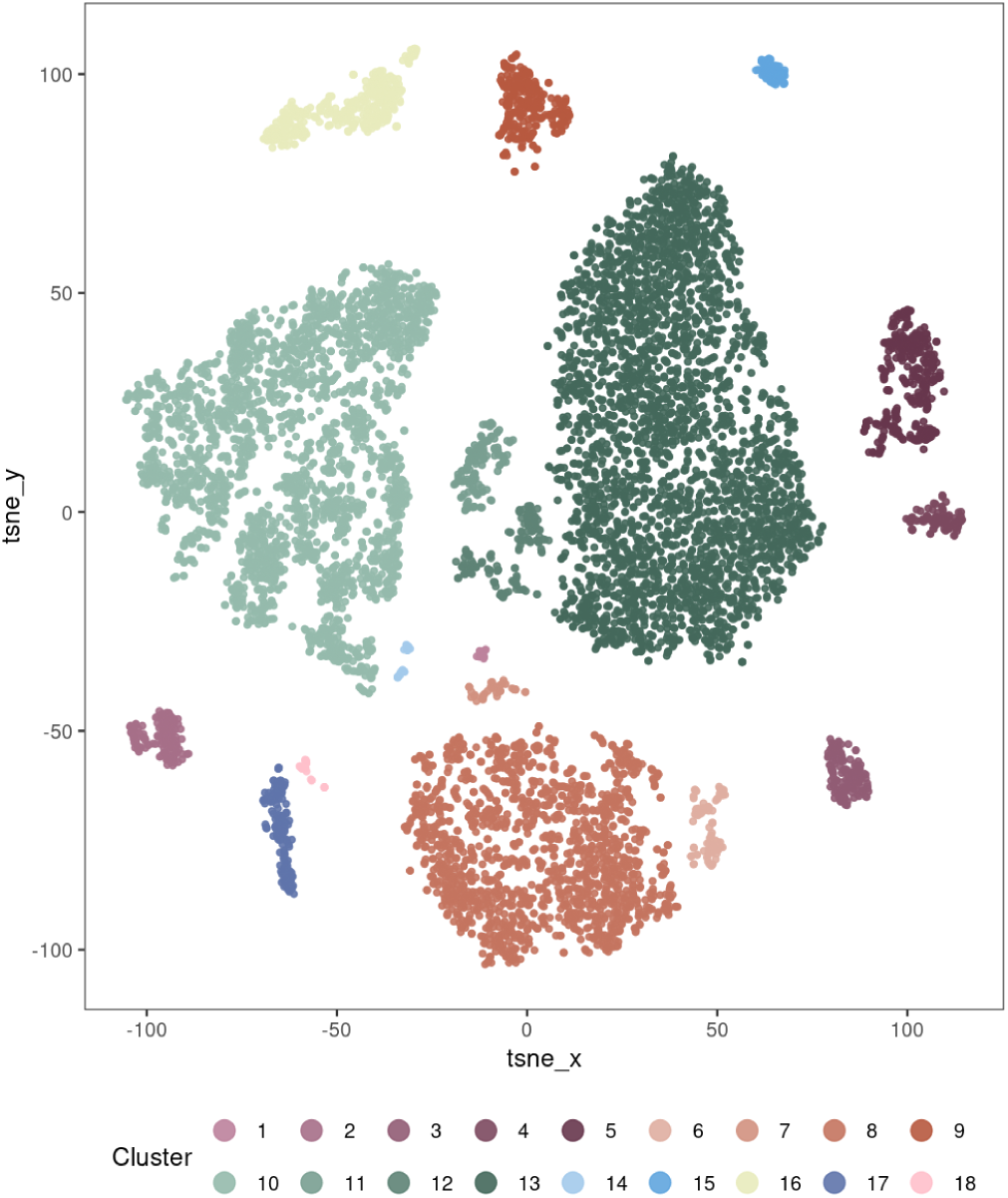
The t-SNE embedding of the ovarian cancer dataset coloured by cluster annotations. The embedding and cluster annotations used were provided by the original paper.

**Supplementary Figure 6.**
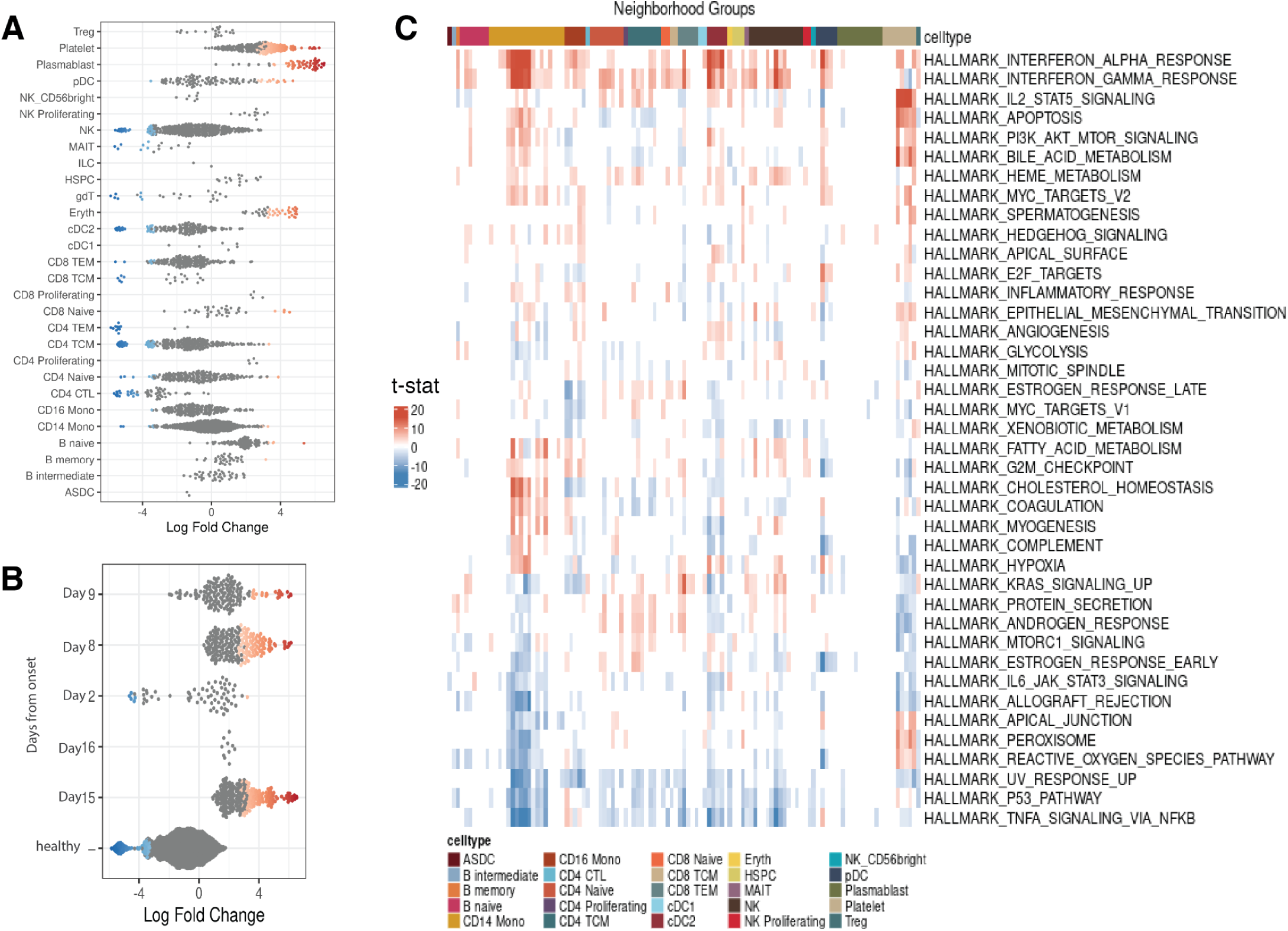
Differential pathway activity test at cell neighborhood level using Milo for Hallmark gene programs selected and quantified by scDECAF. **A.** Differential Abundance (Spatial FDR < 5%) results by Milo summarized at cell type level. **B.** The same results summarized by days from onset. **C.** Heatmap of t-statistics for differential activity of Hallmark pathways selected (among the full Hallmark gene sets) and scored by scDECAF in each Milo neighborhood group, annotated by cell type.

## Description of datasets and processing

**Table 1:** Datasets used for training in each tissue type and corresponding query datasets.

| Tissue type | Dataset type | Description | Access |
| --- | --- | --- | --- |
| PBMC | Query | Sorted healthy donor PBMC | Zheng et al. (via 10x Datasets) |
| PBMC | Reference | 10x donor PBMC | 10x Datasets |
| Dentate Gyrus | Query | Fine-grade postnatal mouse dentate gyrus | Hochgerner et al. (GSE95315) |
| Dentate Gyrus | Reference | Post- and perinatal mouse dentate gyrus | Hochgerner et al. (GSE104323) |
| Pancreas | Query | Pancreatic islets from 4 donors | Baron et al. (via scRNAseq Bioconductor package) |
| Pancreas | Reference | Pancreatic islets from human sample | Segerstolpe et al. (via scRNAseq Bioconductor package) |
| Lung | Query | Lung tissue from mouse | Tabula Muris Consortium |
| Lung | Reference | Mouse lung atlas | Travaglini et al. |

### 10x PBMC datasets

Sorted peripheral blood mononuclear (PBMC) cells from healthy donors from Zheng et al^30^ were used as the “target” or “query” dataset to benchmark reference-based and reference-free cell type annotation methods. Data was subsetted on 5000 HVG for the analysis. Where a single cell reference was required, we used the 8000 PBMCs from a healthy donor^64^ . Data was subsetted on 3000 HVGs for the analysis. Datasets were obtained from the 10x support dataset collection http://support.10xgenomics.com/single-cell/datasets. Data was normalized by SCTransform normalization.

### Pancreas

Human Pancreas datasets from Baron^32^ et al. and Segerstolpe^65^ et al. studies were accessed via the scRNAseq^66^ Biocondutor package. The dataset by Baron et al. contains 10,000 single-cells of pancreatic islets from 4 human donors sequenced by inDrop. This dataset was used as the “target” or “query” dataset to benchmark reference-based and reference-free cell type annotation methods. The dataset by Segerstolpe et al. consists of 2,209 cells from human pancreatic islets. This dataset was used as the reference to annotate cells in Baron et al., where a reference was required to annotate the cells. To run scDECAF and Seurat v4, we used 2000 HVGs and applied SCTransform^28^ normalization. For all other workflows, we applied scran^67^ normalization.

### Dentate Gyrus Brain

Single cells from Peri- and postnatal neurogenesis in the dentate gyrus in mice was obtained from Hochgerner^31^ et al. Transcriptome of 5454 single cells from dentate gyrus sampled at postnatal day 12, 16, 24 and 35 were used as the “target” or “query” for cell type annotation. This dataset is available under GEO accession GSE95315 . Where a reference single cell dataset was required, we used 24185 single cells from dentate gyrus sampled at ages ranging from embryonal day 16.5 to postnatal day 132, available from GSE104323. To run scDECAF and Seurat v4, we used 3000 HVGs and applied SCTransform normalization. For all other workflows, we applied scran normalization.

### Lung

Single cell transcriptomics from 5449 mouse lung cells obtained from Tabula Muris^68^ (Tabula Muris Lung 10x) were used as “target” or “query” for cell type annotation. Where a single cell reference was required, we used 24,618 mouse lung cells from Travaglini^30^ et al. We used 3000 HVGs for the analysis and applied SCTransform normalization.

### Kang PBMC

Single cell transcriptomics of 25,000 PBMC cells from 8 lupus patient samples with and without stimulation with interferon (IFN)-β^21^. Data was downloaded from https://figshare.com/ndownloader/files/34464122. We used 2000 HVGs for the analysis and applied SCTransform normalization.

### Slide-tags snRNA-seq on human prefrontal cortex

Normalized single-nucleus gene expression and spatial coordinates for 4067 cells from the human prefrontal cortex was downloaded from the Broad Institute Single Cell Portal. We used scanpy^69^ to subset the data on 500 (none-spatial) highly variable genes. We then scored the cells for each cell type using top 500 spatially highly variable genes provided by the original study. Predicted cell types were compared to cell type annotations provided by the authors.

**Supplementary note 1.**
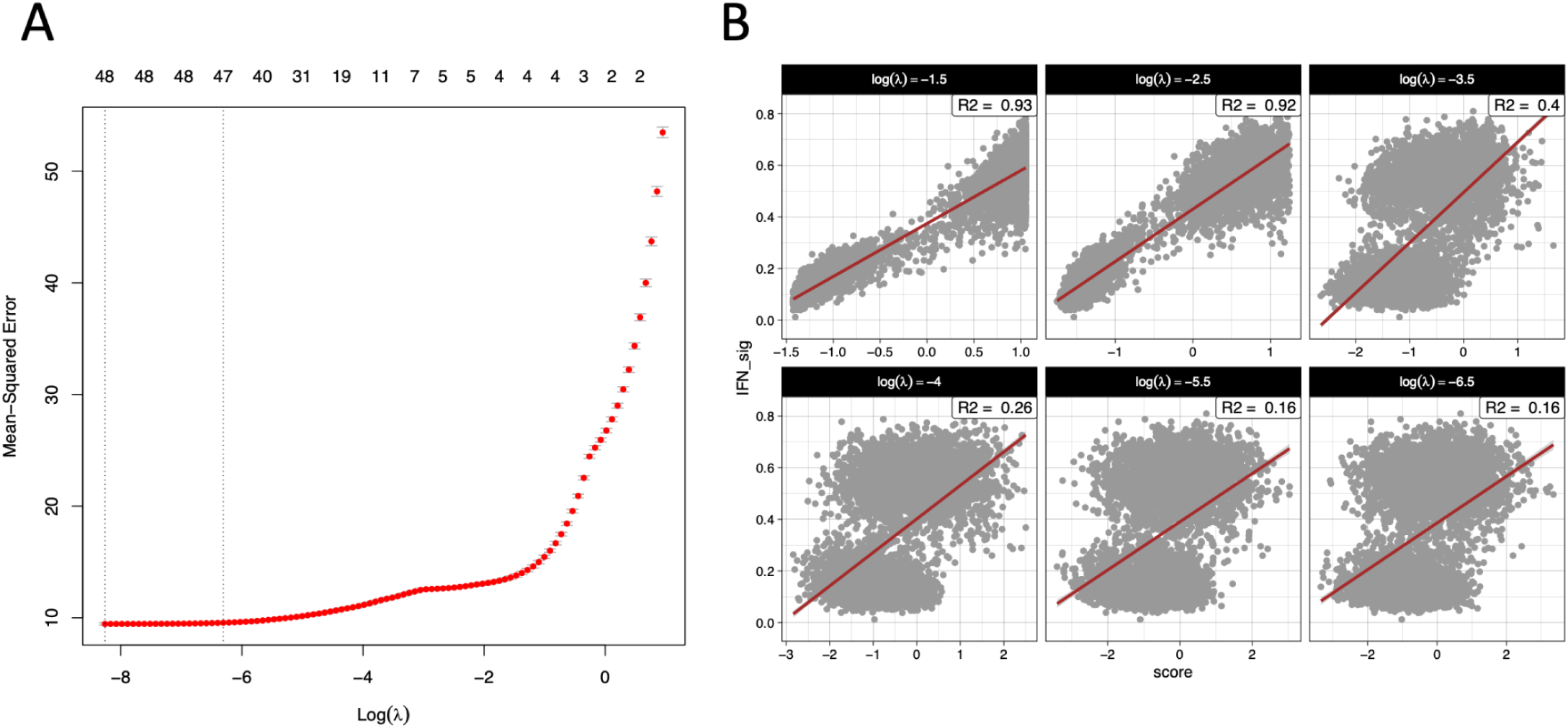
**A.** Example of a sparsity inducing operator plot in Kang et al., IFNbeta-stimulated PBMC dataset. Top is the number of gene sets surviving selection for different values of shrinkage operator lambda (x-axis) and the reconstruction error of the embedding (y-axis). Dotted lines indicate values for lambda which result in least mean squared error for reconstruction of the cell embedding. As the sparsity penalty increases (from left to right), less gene sets are retained, which is also coupled with highest mean squared error in the reconstruction of cell embedding. **B.** Adjusted R^2^ for the association between scDECAF scores (x-axis) and IFNalpha signature scores (y-axis) in Monocyte cells in Kang et al. for different values of shrinkage operator, lambda. The association between scDECAF scores and IFN signature of cells decreases as more gene sets are included in the set of input gene sets. This highlights the importance of using the sparsity selection mode prior to learning vector representations of the gene sets.

